# First observation of *Aedes albopictus* in the Tshuapa province (Boende) of the Democratic Republic of the Congo

**DOI:** 10.1101/2021.08.03.454759

**Authors:** Joachim Mariën, Nicolas Laurent, Nathalie Smitz, Sophie Gombeer

**Author notes:** **Corresponding author:** Joachim Mariën Universiteitsplein 1, 2610 Antwerpen, Belgium.

## Abstract

In May-June 2021, we detected *Aedes albopictus* adults near the central hospital in Boende, the capital city of the Tshuapa province of the Democratic Republic of the Congo (DRC). We identified the mosquitoes using morphological and molecular techniques (COI barcoding). This is the first report of this species in the DRC outside of Kinshasa and Kongo Central. Given the central location of Boende in the Congo Basin, our finding suggests that the vector might also have spread to other cities which are located upstream of the Congo River and its major tributaries. Because *Aedes albopictus* is an important vector for human arbovirus transmission, we highlight the need to investigate its distribution range and to update disease risk maps in Central Africa.

## Introduction

*Aedes albopictus* (Skuse 1895) (Diptera: Culicidae) is considered to be the most invasive mosquito species worldwide^1^. The species originated in Southeast Asia and has successfully expanded to other parts of the world during the last decades^2^. This range expansion is primarily caused by human activities related to global commerce (e.g. trafficking of used tires) and the mosquito’s adaptation to various environments in both tropical and temperate regions (e.g. ability to breed in different container forms)^3–5^. This rapid expansion has also been linked to multiple outbreaks of *Aedes*-borne disease, such as Chikungunya (CHIKV), Zika, yellow fever and Dengue^6–8^.

Although the first observation of *Ae. albopictus* in Africa was reported four decades ago (Nigeria), its presence in the Democratic Republic of Congo (DRC) was only described for the first time in 2018^9^. One year later, we linked a major CHIKV outbreak to its recent colonization in that area (the provinces Kinshasa and Kongo Central during 2019-2020)^10,11^. Indeed, *Ae. albopictus* was the main *Aedes* species around human CHIKV cases in Kinshasa (>80% of the adults and larvae), Kasangulu (>99% of the adults) and Matadi (>95% of the adults and larvae). The only other *Aedes* species detected during this outbreak was *Aedes aegypti,* a primary vector of arboviruses indigenous to Africa. Furthermore, we could link the CHIKV outbreak to an amino acid substitution in the viral envelope gene E1 (E1-A226V) which makes the virus more suitable for dissemination by *Ae. albopictus*^10^. These previous results indicate that *Ae. albopictus* has now well-established populations in South-Western DRC, highlighting the potential for future outbreaks of *Aedes*-borne diseases and the urgent need for additional surveillance and control methods in the entire country^11,12^.

To prevent and control future outbreaks of *Aedes*-borne viruses in the Congo Basin, it will be crucial to monitor both the viruses and their vectors in the region^13^. While research in the DRC is often restricted to Kinshasa due to logistic reasons, the eco-epidemiological drivers of disease transmission could be significantly different in other regions (e.g. through differences in seasonal dynamics, insecticide resistance, landscape elements, human-social behaviours, immunological responses, etc…). Here, we provide data on the presence of *Ae. albopictus* in Boende to encourage scientists to investigate the ecology of *Ae. albopictus* upstream of the Congo River and warn policymakers for a potential increase of *Aedes*-borne viral outbreaks in the area.

## Methods

In May 2021, we were bitten by mosquitoes while walking in the garden of our hotel in Boende (Tshuapa province, 0°16’40.4”S and 20°52’39’’E). We identified these mosquitoes as *Ae. albopictus* by their characteristic white stripe on the scutum. Six female *Ae. albopictus* were collected during human-landing and confirmed morphologically following Walter Reed’s identification keys^14^. Later that day we went to the central hospital in Boende (0°16’38.5’’S 20°52’59’’E) and were again bitten by the same species. One month later (June 2021), *Ae. albopictus* mosquitoes bit us again when we returned to the same two locations.

The morphological identifications were validated by DNA-barcoding, a technique based on the amplification of the partial mitochondrial cytochrome c oxidase subunit I (COI) gene^15^. Sanger sequencing of the 658-base pair barcode and phylogenetic analyses followed laboratory protocols and methodologies described in Wat’senga Tezzo et al 2021^16^. In short, PCR amplicons (EPPO 2016; LCO1490 and HCO2198 universal primers) and negative controls were checked on an agarose gel, purified and sequenced in both directions on an ABI 3230xl capillary DNA sequencer using BigDye Terminator v3.1 chemistry (ThermoFisher Scientific). Generated sequences were processed using Geneious® R11 (Biomatters Ltd., Auckland, New Zealand) for quality control, pairing of bi-directional strands, and to extract one consensus sequence per specimen. The generated consensus sequences (N_Boende_ = 5) were compared against the BOLD Identification System with Species Level Barcode Records.

To check the phylogenetic relation of the sequences, we compiled a list of *Aedes* species (belonging to the genus *Stegomyia*) that are considered to be of medical importance in the Afrotropical region by consulting the WRBU and the APHC databases^16^. Publically available COI sequences from these species were cleaned and aligned using ClustalW in Geneious® R11. *Culex quinquefasciatus* was added as outgroup to the alignment. A Neighbour-joining tree was constructed assuming the Kimura 2-paramers and branch support assessed by 500 bootstrap (BS) replicates using MEGA7 ^17–19^. The generated sequences were deposited in GenBank with following accession numbers: MZ673312-MZ673316.

## Results and Discussion

The observed *Ae. Albopictus* during the months of May and June 2021 in Boende suggest that established populations of the mosquito species occur close to the Tshuapa river in Boende. All *Ae. albopictus* COI sequences including the newly generated sequences group together in a highly supported cluster (100 BS, **Figure 1**), validating the morphological identifications. Indeed, all similarity percentages reached 100% to *Ae. Albopictus* in the BOLD identification system, with the identification of one haplotype in Boende that also occurred in Kinshasa.

**Figure 1.**
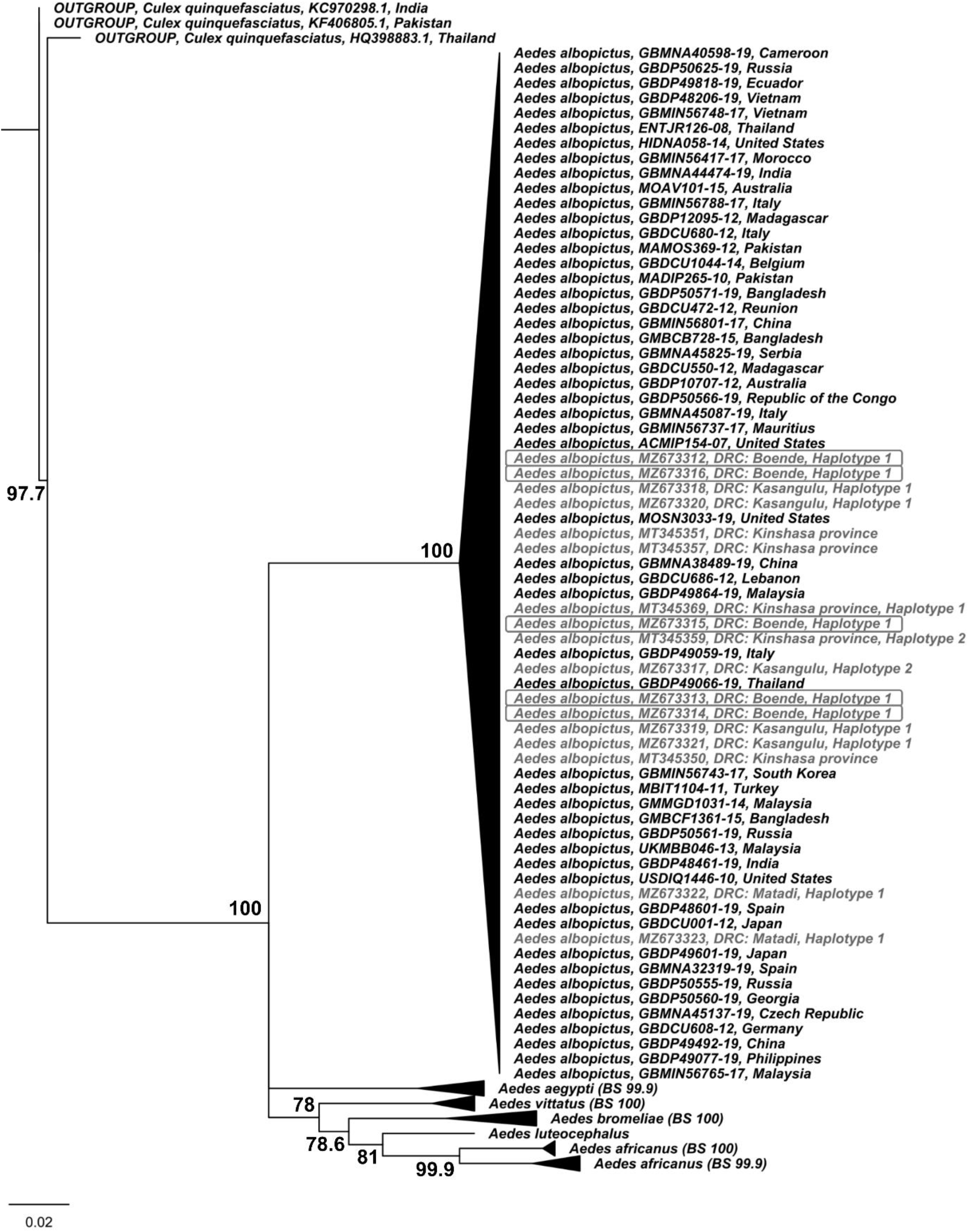
Neighbor-joining tree (1,000 bootstrap, 75 support threshold) including the six medically important *Aedes* species of the subgenus *Stegomyia* occurring in the Afrotropical region. Available COI sequences of *Aedes albopictus* of the Democratic Republic of the Congo (DRC) are highlighted in grey including the sequences generated in this study from Boende (in boxes).

Although Boende is situated more than 750km from Kinshasa in the middle of the rain forest, the finding of *Ae. albopictus* in the city is not unexpected^13^. Boende is located on the left bank of the Tshuapa river (one of the tributaries of the Congo River) and boats arrive daily from downstream large cities such as Mbandaka and Kinshasa. These boats are usually very crowded and stacked with various goods, including water containers and used tires, making them ideal vehicles for *Ae. albopictus* larvae and eggs^20^. This assumption is further supported by the fact that the identified haplotype in Boende is shared with ones recorded in Kinshasa and Kongo Central (**Figure 1**)^16^, also being the most frequent in the latter study (Haplotype_frequency_ = 0.71). Furthermore, the average precipitation (210.9 cm per year) and daily temperatures (ranging between 24°C and 30°C year-round) fall within the optimal boundaries for *Ae. albopictus* development^8,21^. Given that many cities in the DRC have similar climate conditions and transportation links to Kinshasa, we expect that established populations of *Ae. albopictus* also exist in other cities located near the Congo River or its tributaries throughout the Congo Basin. If this is the case, we predict that the frequency and magnitude of *Aedes*-borne viral outbreaks will increase significantly the following years across Central DRC, similar to what we currently observe in South-Western DRC^10^.

Further research is clearly necessary to investigate *Ae. albopictus’* distribution and persistence (established or transient populations) in Boende and the rest of the country.

## Ethics statement

Ethical approval is not required for this type of study.

## Acknowledgements

This research was supported through the BIODIV-AFREID project (BiodivERsA programme, www.biodiversa.org, with funding organization FWO) and the EBOVAC3 project. We are grateful for the logistic support of the University of Kinshasa, the University of Kisangani and the Ebovac3 team from the University of Antwerp.

## Availability of data and materials

The biological material and data used in the current study are available from the corresponding author on reasonable request. The generated COI sequences were deposited in GenBank with following accession numbers: MZ673312-MZ673316.

